# Collagen-dependent DDR1 ectodomain shedding regulates proliferative signaling in radiation-resistant glioblastoma

**DOI:** 10.64898/2026.10.08.757755

**Authors:** Sophia K. Dunlap, Caitlin A. Harvey, Amr Elkholy, Leah Gehrs, Mostafa Mohamed, Saeed Zakakhosravi, Taylor L. Schanel, James M. Murphy, Ssang-Taek Steve Lim, Christopher D. Willey, Eun-Young Erin Ahn

## Abstract

Glioblastoma (GBM) is characterized by near-universal recurrence and resistance to therapy. Although tumor-intrinsic mechanisms of radiation resistance have been extensively studied, how the extracellular matrix influences signaling in radiation-resistant GBM remains poorly understood. Here, we investigated discoidin domain receptor 1 (DDR1), a collagen-activated receptor tyrosine kinase, using matched radiation-sensitive parental and radiation-resistant patient-derived GBM models generated through serial *in vivo* irradiation. Within a three-dimensional collagen I matrix, radiation-resistant GBM cells exhibited distinct temporal patterns of Ki-67 positivity and ERK1/2 phosphorylation compared with parental cells. Despite increased DDR1 expression, radiation-resistant cells exhibited markedly reduced DDR1 phosphorylation and enhanced ectodomain shedding, characterized by accumulation of the membrane-associated C-terminal fragment and release of the N-terminal fragment. DDR1 depletion reduced ERK1/2 phosphorylation, supporting its contribution to ERK1/2 signaling under these conditions. ADAMTS1 was markedly upregulated in radiation-resistant cells, and its knockdown reduced DDR1 ectodomain shedding while increasing full-length DDR1 abundance, ERK1/2 and RB phosphorylation, cyclin D1 expression, and Ki-67 positivity. Together, these findings reveal a distinct DDR1 regulatory state in acquired radiation-resistant GBM, characterized by enhanced ADAMTS1-associated ectodomain shedding, reduced receptor phosphorylation, and attenuated proliferative signaling within a three-dimensional collagen I matrix.

## INTRODUCTION

Glioblastoma (GBM), IDH-wildtype, is the most aggressive primary brain tumor and is classified as a WHO grade 4 diffuse astrocytic glioma (1). Despite standard-of-care treatment with maximal safe resection, radiation therapy, and temozolomide chemotherapy, the 5-year survival rate is less than 7%, with a median survival of 12-15 months following diagnosis (2, 3). Treatment failure is largely driven by the near-inevitable recurrence of GBM following initial therapy (4). Several factors contribute to the high recurrence rate. GBM is a highly invasive tumor with poorly defined borders, limiting the ability of surgical resection to eliminate all malignant cells. Consequently, infiltrating tumor cells remain within the surrounding brain tissue and can contribute to subsequent tumor recurrence (5). Treatment resistance represents another major contributor to GBM recurrence and treatment failure. Although MGMT promoter methylation is an established predictor of response to temozolomide, radiation resistance remains difficult to predict and is not fully understood (6, 7). Additionally, key factors hamper the development of novel GBM treatments, including the blood-brain barrier, which limits drug uptake; tumor heterogeneity, which allows subpopulations to survive and repopulate; and an immune-cold microenvironment that limits the efficacy of immunotherapy (8–11). Collectively, these challenges underscore the need to better understand the mechanisms by which GBM cells acquire and maintain resistance to radiation therapy. Although tumor-intrinsic molecular mechanisms of radiation resistance have been widely investigated, the contribution of the tumor microenvironment, particularly the extracellular matrix (ECM), to the behavior and signaling of radiation-resistant GBM cells remains incompletely understood.

The ECM comprises three main components: fibrous proteins such as elastin and collagens, proteoglycans and glycosaminoglycans, and glycoproteins including laminin and fibronectin (12). Once viewed as a structural scaffold, the ECM is now recognized as a dynamic regulator of tumor behavior and therapy resistance, not only by providing structural support and tissue integrity, but also by transmitting biochemical signals essential for cellular regulation (12). In radiation resistance, several ECM-related factors are altered, including matrix stiffness and collagen deposition (13–15). Compared with normal brain tissue, GBM exhibits increased expression and remodeling of collagens and altered activity of matrix metalloproteases (16–19). Collagen-binding receptors provide a critical interface through which these changes in the extracellular environment are sensed and translated into intracellular signaling programs in tumor cells.

One collagen-specific receptor, discoidin domain receptor 1 (DDR1), has been implicated in therapy resistance mechanisms (20). DDR1 is a receptor tyrosine kinase (RTK) that is activated by several types of collagen, including collagens I-V (21, 22). DDR1 contains structural and biochemical features required for ligand recognition and activation, including an extracellular discoidin domain containing collagen-binding sites and a transmembrane leucine zipper that facilitates receptor association. (23–25). DDR1 exhibits distinctive activation kinetics: collagen binding promotes receptor clustering and kinase activation, resulting in receptor autophosphorylation (26, 27). DDR1 phosphorylation includes Y792 within the kinase domain and Y513 within the intracellular juxtamembrane region (26–28). Unlike the rapid and transient activation characteristic of many RTKs, collagen-induced DDR1 phosphorylation develops slowly and can remain sustained for prolonged periods (29).

DDR1 phosphorylation and signaling are further regulated by ectodomain shedding and endocytosis. Ectodomain shedding involves proteolytic cleavage and release of the extracellular N-terminal fragment, leaving a membrane-associated C-terminal fragment, and can attenuate receptor phosphorylation and signaling (29–33). Recent studies have also identified functional roles for the shed N-terminal DDR1 ectodomain, including the regulation of collagen fiber alignment and immune exclusion in breast cancer (34, 35). Despite these established mechanisms of DDR1 regulation, whether DDR1 signaling and ectodomain shedding are altered in radiation-resistant GBM remains unclear.

Although DDR1 has been studied in GBM, how DDR1 expression, activation, and proteolytic processing are altered in radiation-resistant GBM within a three-dimensional collagen I environment remains poorly understood. Moreover, the regulation of DDR1 in matched radiation-sensitive and acquired radiation-resistant patient-derived GBM models has not been defined.

Although DDR1 ectodomain shedding and its effects on receptor signaling have been described in other cellular systems, whether DDR1 processing is altered during the acquisition of radiation resistance in GBM remains unknown. Here, we investigated DDR1 regulation in matched radiation-sensitive parental and acquired radiation-resistant patient-derived GBM models cultured in a three-dimensional collagen I matrix. We examined differences in DDR1 expression, ectodomain shedding, receptor phosphorylation, and proliferative signaling and investigated the contribution of ADAMTS1 to DDR1 processing. Our findings reveal a distinct DDR1 regulatory state in radiation-resistant GBM, characterized by increased receptor abundance but enhanced ectodomain shedding and reduced phosphorylation, accompanied by altered proliferative signaling.

## RESULTS

### Radiation-resistant GBM cells exhibit distinct temporal patterns of proliferative activity and ERK1/2 signaling in a 3D collagen I matrix

To assess collagen I expression in GBM, we analyzed COL1A1 and COL1A2 mRNA and protein levels using UALCAN. Both COL1A1 and COL1A2 mRNA levels were significantly increased in GBM compared with normal brain tissue, whereas only COL1A2 protein levels were significantly elevated (**Figure 1A, B**). To investigate acquired radiation resistance, we utilized matched radiation-sensitive parental (P) and radiation-resistant (RT) JX39 GBM patient-derived xenograft (PDX) models. The radiation-resistant JX39P-RT model was generated from the parental PDX through serial *in vivo* irradiation (2 Gy per fraction, six fractions per round, for 6–8 rounds), as previously described by Stackhouse et al. (36) (**Figure 1C**).

**Figure 1.**
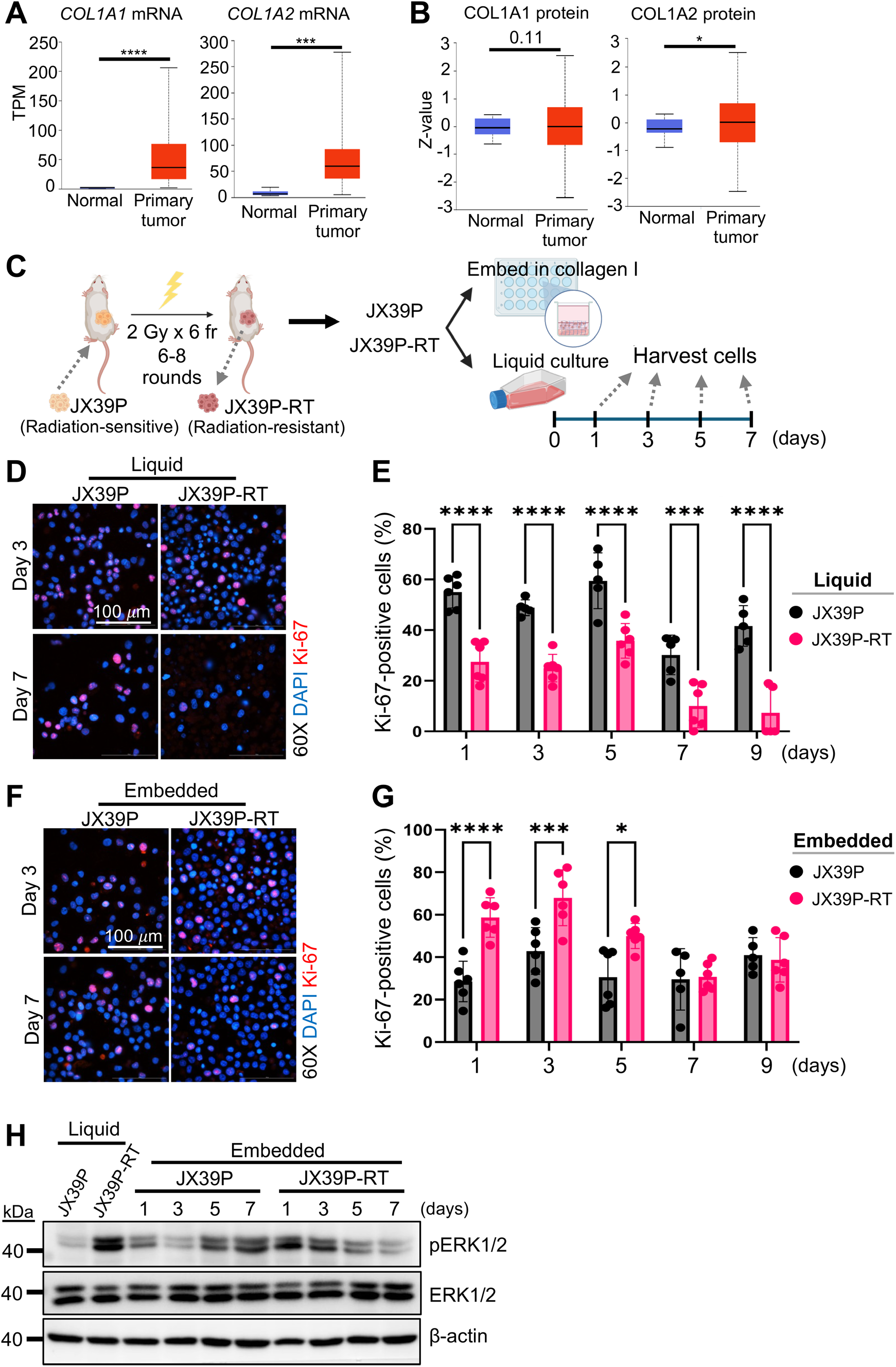
Radiation-sensitive and radiation-resistant GBM cells exhibit distinct temporal patterns of proliferative activity and ERK1/2 signaling in a three-dimensional collagen I matrix. (A) Analysis of TCGA data using UALCAN showing significantly increased *COL1A1* and *COL1A2* mRNA expression in GBM (n = 156) compared with normal brain tissue (n = 5). *COL1A1*, \*\*\*\**p* = 4.69 × 10⁻⁶; *COL1A2*, \*\*\**p*= 1.2 × 10⁻⁴. (B) Analysis of CPTAC proteomic data using UALCAN showing significantly increased COL1A2 protein expression, but no significant difference in COL1A1 protein expression, in GBM (n = 18) compared with normal brain tissue (n = 199). COL1A2, \**p* = 0.0208; COL1A1, *p* = 0.105. (C) Schematic illustrating the generation of paired radiation-sensitive parental (JX39P) and radiation-resistant (JX39P-RT) GBM patient-derived xenograft (PDX) models through serial *in vivo* irradiation (2 Gy per fraction, six fractions per round, for 6–8 rounds). JX39P and JX39P-RT cells were cultured in liquid suspension or embedded in a three-dimensional collagen I matrix and harvested at the indicated time points (days 1, 3, 5, and 7). (D) Representative immunofluorescence images of Ki-67 (red) and DAPI (blue) staining in JX39P and JX39P-RT cells cultured in liquid suspension. Scale bars, 100 µm. (E) Quantification of Ki-67-positive cells in liquid suspension cultures, expressed as the percentage of total DAPI-positive nuclei. Radiation-sensitive JX39P cells exhibited a higher proportion of Ki-67-positive cells than JX39P-RT cells. \*\*\**p*<0.001, \*\*\*\**p*<0.0001. (F) Representative immunofluorescence images of Ki-67 (red) and DAPI (blue) staining in JX39P and JX39P-RT cells embedded in collagen I for the indicated durations. Scale bars, 100 µm. (G) Quantification of Ki-67-positive cells in collagen I-embedded cultures at the indicated time points, expressed as the percentage of total DAPI-positive nuclei. JX39P-RT cells exhibited higher Ki-67 positivity at early time points, followed by a decline at later time points. \**p*<0.05, \*\*\**p*<0.001, \*\*\*\**p*<0.0001. (H) Western blot analysis of phosphorylated ERK1/2 (pERK1/2) and total ERK1/2 in JX39P and JX39P-RT cells cultured in liquid suspension or embedded in collagen I for the indicated durations, showing distinct temporal patterns of ERK1/2 phosphorylation between parental and radiation-resistant cells.

To determine whether a 3D collagen I matrix differentially influences proliferative activity in radiation-sensitive and radiation-resistant GBM cells, JX39P and JX39P-RT cells were embedded in a 3D collagen I matrix or maintained in liquid suspension without collagen I as controls. Cells were harvested at days 1, 3, 5, and 7 after culture initiation (**Figure 1C**). Proliferative activity was assessed by immunofluorescence staining for Ki-67, with DAPI used to visualize nuclei. In liquid suspension, radiation-sensitive JX39P cells exhibited a higher percentage of Ki-67-positive cells than radiation-resistant JX39P-RT cells (**Figure 1D, E**). In contrast, following embedding in the 3D collagen I matrix, JX39P-RT cells exhibited increased Ki-67 positivity at early time points (days 1 and 3), followed by a decline at later time points (**Figure 1F, G**). These findings indicate that radiation-resistant GBM cells exhibit a distinct, time-dependent pattern of proliferative activity within the 3D collagen I matrix.

Because MAPK signaling activation through ERK1/2 is an established regulator of cell proliferation (37, 38), we next examined ERK1/2 phosphorylation under liquid suspension and 3D collagen I culture conditions. Under liquid suspension conditions, JX39P and JX39P-RT cells exhibited relatively similar basal levels of ERK1/2 phosphorylation. However, following embedding in the 3D collagen I matrix, the two cell populations displayed distinct temporal patterns of ERK1/2 phosphorylation (**Figure 1H**). In radiation-sensitive JX39P cells, ERK1/2 phosphorylation was initially low but progressively increased over time, whereas JX39P-RT cells exhibited a different temporal phosphorylation pattern. Together, these findings demonstrate that radiation-sensitive and radiation-resistant GBM cells exhibit distinct temporal patterns of Ki-67 positivity and ERK1/2 activation when cultured in a 3D collagen I matrix.

### DDR1 is upregulated at both the mRNA and protein levels and regulates ERK1/2 signaling in radiation-resistant GBM

Given the distinct proliferative and ERK1/2 signaling responses of JX39P-RT cells to collagen I, we next investigated DDR1, a collagen-activated receptor tyrosine kinase with established roles in cell proliferation and MAPK/ERK signaling (28, 39). First, to assess DDR1 expression in GBM, we analyzed DDR1 mRNA and protein levels using UALCAN and found increased DDR1 expression in GBM compared with normal brain tissue (**Figure 2A**, **B**).

**Figure 2.**
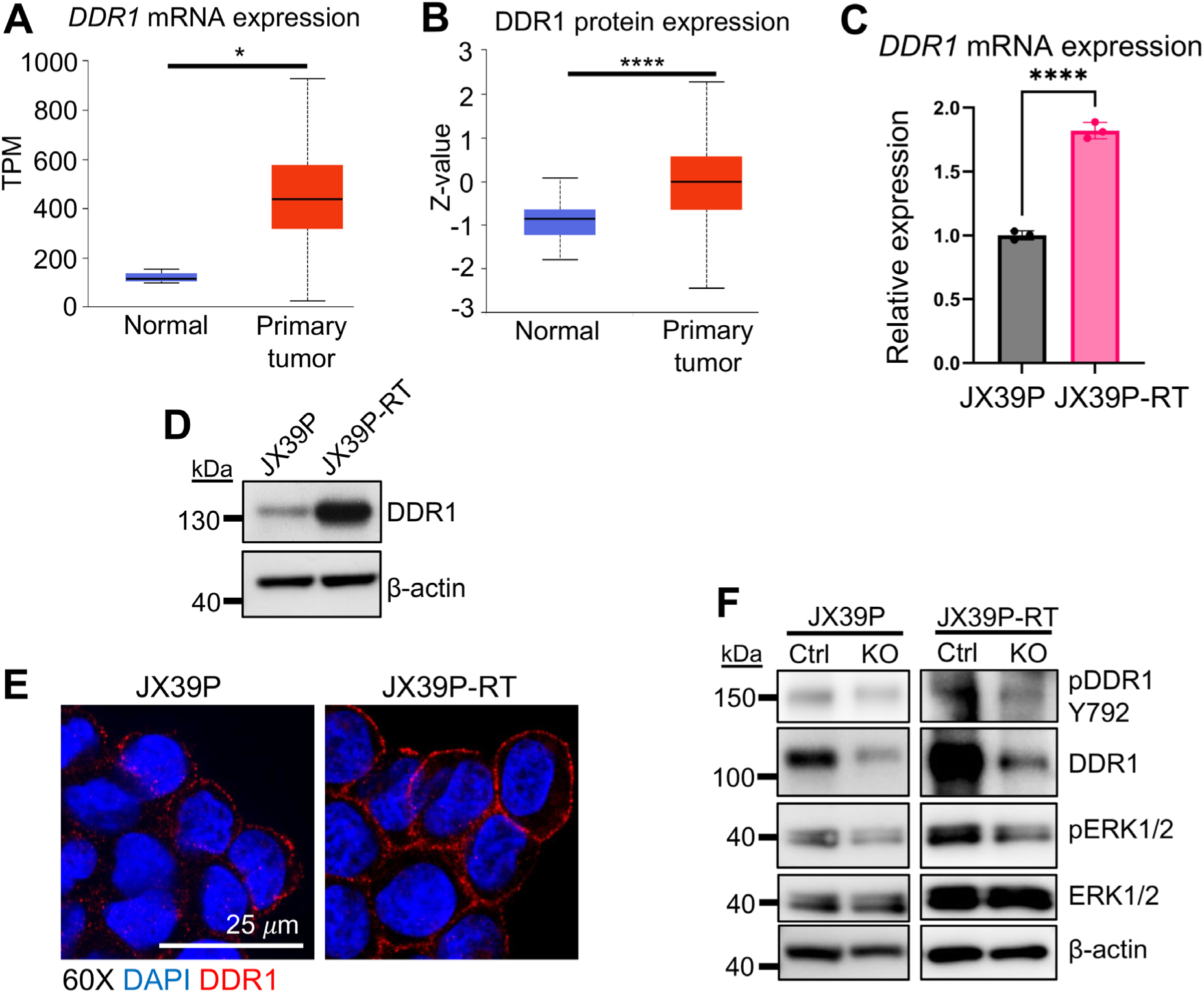
DDR1 is upregulated in radiation-resistant GBM and contributes to ERK1/2 activation in a collagen I environment. **(A)** Analysis of TCGA data using UALCAN showing significantly increased *DDR1* mRNA expression in GBM (n = 156) compared with normal brain tissue (n = 5). \**p* = 0.0208. **(B)** Analysis of CPTAC proteomic data using UALCAN showing significantly increased DDR1 protein expression in GBM (n = 199) compared with normal brain tissue (n = 18). \*\*\*\**p* = 1.11 × 10⁻¹¹. **(C)** Quantitative RT-PCR analysis of *DDR1* mRNA expression in radiation-sensitive JX39P and radiation-resistant JX39P-RT cells, showing increased expression in JX39P-RT cells. \*\*\*\**p*<0.0001. **(D)** Western blot analysis of DDR1 protein expression in JX39P and JX39P-RT cells. **(E)** Representative confocal immunofluorescence images of DDR1 (red) and DAPI (blue) staining in JX39P and JX39P-RT cells, showing increased DDR1 expression and membrane-associated localization in JX39P-RT cells. Scale bar, 25 µm. **(F)** Western blot analysis of DDR1, phosphorylated DDR1 (pY792), phosphorylated ERK1/2 (pERK1/2), and total ERK1/2 in control and CRISPR/Cas9-mediated DDR1 knockout (KO) JX39P and JX39P-RT cells following 24 h of collagen I embedding.

*In vitro* analysis of JX39P and JX39P-RT cells revealed increased DDR1 mRNA and protein levels in radiation-resistant cells compared with their radiation-sensitive parental counterparts, as determined by qRT-PCR and Western blot analysis (**Figure 2C, D**). Consistent with these data, confocal immunofluorescence (IF) analysis of JX39P/RT cells demonstrated increased DDR1 expression in radiation-resistant GBM and confirmed its localization at the cell membrane (**Figure 2E**).

To determine whether DDR1 contributes to ERK1/2 signaling in the collagen I environment, we generated DDR1 knockout (KO) JX39P and JX39P-RT cells using CRISPR/Cas9. Control and DDR1 KO JX39P/RT cells were embedded in collagen I for 1 day and harvested. DDR1 loss was confirmed by Western blot analysis and was accompanied by decreased phosphorylation at Y792. Compared with control cells, DDR1 KO cells also exhibited reduced phosphorylation of ERK1/2 (**Figure 2F**), supporting a role of DDR1 in ERK1/2 activation in collagen I-embedded GBM cells, consistent with previous reports (28, 39).

These findings reveal increased DDR1 abundance in acquired radiation-resistant GBM cells and support a contribution of DDR1 to ERK1/2 signaling within a 3D collagen I matrix, prompting us to investigate whether DDR1 phosphorylation and proteolytic processing are also altered in the resistant state.

### DDR1 ectodomain shedding is enhanced in radiation-resistant GBM cells embedded in a 3D collagen I matrix

To investigate DDR1 processing in radiation-resistant GBM cells embedded in a 3D collagen I matrix, JX39P/RT cells were either embedded in collagen I or cultured in liquid suspension and harvested at multiple time points. Western blot analysis using an antibody recognizing the intracellular C-terminal region of DDR1 revealed the accumulation of a lower-molecular-weight DDR1 fragment consistent with the membrane-associated C-terminal fragment (CTF) generated following ectodomain shedding. In contrast, analysis with an antibody recognizing the DDR1 extracellular domain showed reduced detection of full-length DDR1 in radiation-resistant JX39P-RT cells compared with parental JX39P cells (**Figure 3A, B**). Together, these findings were consistent with enhanced DDR1 ectodomain shedding in JX39P-RT cells within the collagen I environment.

**Figure 3.**
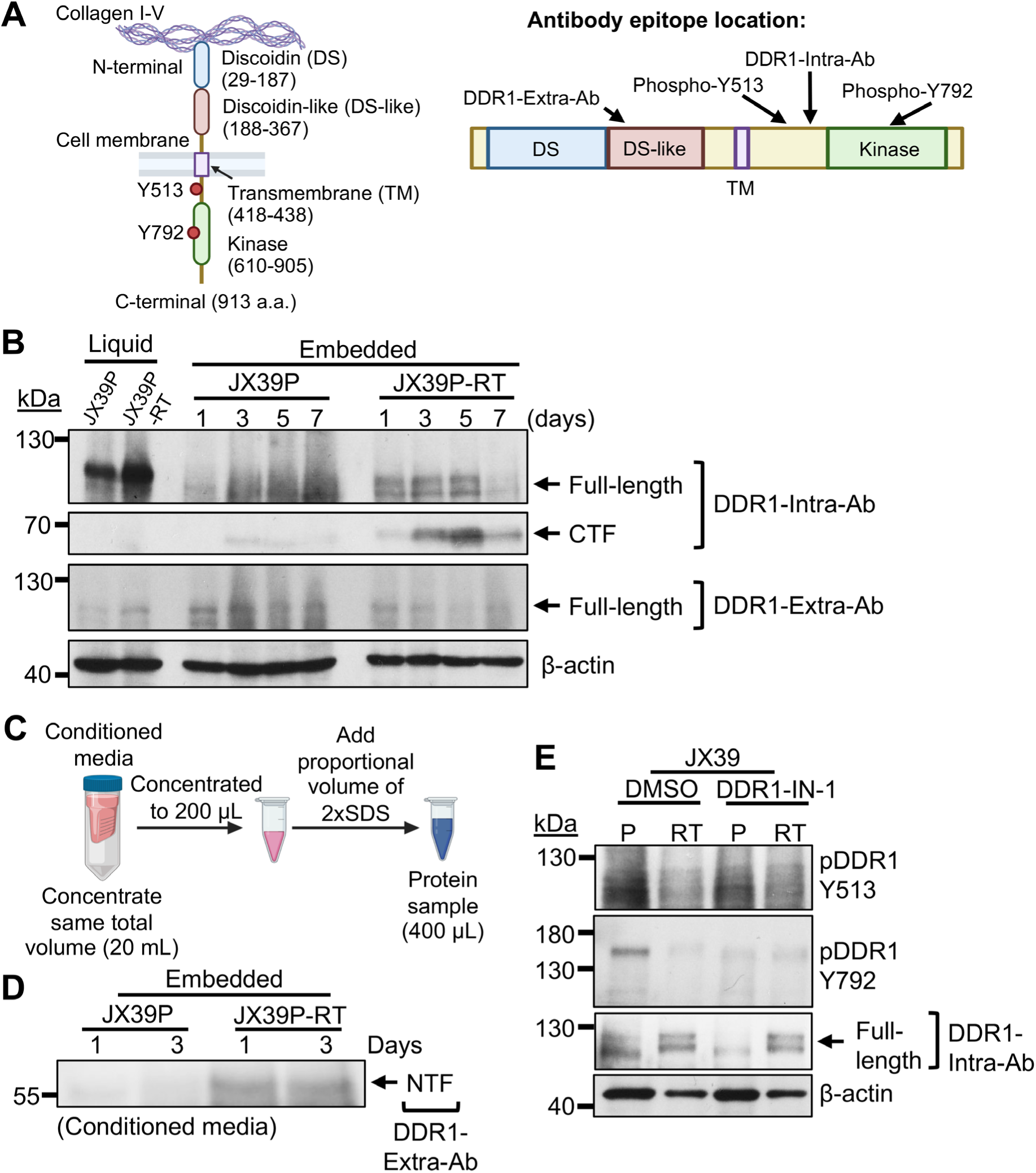
Radiation-resistant GBM cells exhibit enhanced DDR1 ectodomain shedding and reduced DDR1 phosphorylation in a collagen I environment. **(A)** Schematic illustrating DDR1 cell membrane localization, domain organization, and antibody recognition sites. An antibody recognizing the extracellular domain of DDR1 (DDR1-Extra-Ab) was used to detect full-length DDR1 and the shed N-terminal fragment (NTF), whereas an antibody recognizing the intracellular domain (DDR1-Intra-Ab) was used to detect full-length DDR1 and the membrane-associated C-terminal fragment (CTF). Phospho-specific antibodies recognizing DDR1 phosphorylated at Y513 and Y792 were used to assess receptor phosphorylation. **(B)** Western blot analysis of JX39P and JX39P-RT cell lysates collected from liquid suspension or collagen I-embedded cultures at the indicated time points using antibodies recognizing the intracellular and extracellular domains of DDR1. Increased accumulation of the DDR1 CTF and reduced detection of extracellular-domain-containing full-length DDR1 were observed in JX39P-RT cells following collagen I embedding. **(C)** Schematic illustrating the preparation of conditioned medium for detection of shed DDR1 fragments. Conditioned medium (20 mL per sample) was concentrated to 200 µL before preparation for Western blot analysis. **(D)** Western blot analysis of concentrated conditioned medium from collagen I-embedded JX39P and JX39P-RT cells using the DDR1 extracellular-domain antibody, showing increased release of the shed DDR1 NTF in JX39P-RT cells. **(E)** Western blot analysis of DDR1 phosphorylation at Y513 and Y792 in JX39P and JX39P-RT cells embedded in collagen I and treated with vehicle (DMSO) or the DDR1 kinase inhibitor DDR1-IN-1 (10 µM). DDR1 phosphorylation was reduced following DDR1-IN-1 treatment in parental JX39P cells, whereas JX39P-RT cells exhibited markedly lower basal DDR1 phosphorylation with little apparent additional reduction following inhibitor treatment.

To further assess DDR1 ectodomain shedding, conditioned media were collected from the collagen I-embedded cultures and concentrated for immunoblot analysis using the extracellular-domain DDR1 antibody (**Figure 3C**). Increased levels of the shed N-terminal fragment (NTF) were detected in conditioned media from radiation-resistant JX39P-RT cells (**Figure 3A, D**), providing additional evidence of enhanced DDR1 ectodomain shedding in radiation-resistant GBM cells.

We next examined whether DDR1 phosphorylation also differed between radiation-sensitive and radiation-resistant GBM cells in the collagen I environment. JX39P and JX39P-RT cells were embedded in collagen I and treated with the DDR1-specific kinase inhibitor DDR1-IN-1 (10 µM). DDR1 phosphorylation at Y792 and Y513 was readily detected in radiation-sensitive parental JX39P cells treated with DMSO and was reduced following DDR1-IN-1 treatment, consistent with DDR1 kinase activity under these conditions. In contrast, markedly lower levels of DDR1 phosphorylation were detected in radiation-resistant JX39P-RT cells, and DDR1-IN-1 treatment had little apparent effect (**Figure 3E**).

Together, these findings reveal a distinct pattern of DDR1 regulation in acquired radiation-resistant GBM cells, characterized by increased DDR1 expression despite enhanced ectodomain shedding and markedly reduced receptor phosphorylation within a 3D collagen I matrix. This contrasting relationship between DDR1 abundance and activation suggests that altered receptor processing is an important feature of the radiation-resistant GBM state.

### Broad metalloprotease inhibition suppresses DDR1 ectodomain shedding but does not enhance proliferation in radiation-resistant GBM

Matrix metalloproteases (MMPs) and a disintegrin and metalloprotease 10 (ADAM10) have been implicated in DDR1 ectodomain cleavage and shedding (31–33); however, the proteases involved in DDR1 ectodomain shedding in GBM, particularly in the context of radiation resistance, remain unclear. To determine whether metalloprotease inhibition affects DDR1 ectodomain shedding, downstream signaling, and proliferation in radiation-resistant GBM, JX39P/RT cells were embedded in collagen I and treated with 10 µM Marimastat (MST), a broad metalloprotease inhibitor. Analysis of conditioned media showed reduced levels of the shed DDR1 NTF following MST treatment in JX39P-RT cells, demonstrating that broad metalloprotease inhibition suppresses DDR1 ectodomain shedding under these conditions (**Figure 4A**).

**Figure 4.**
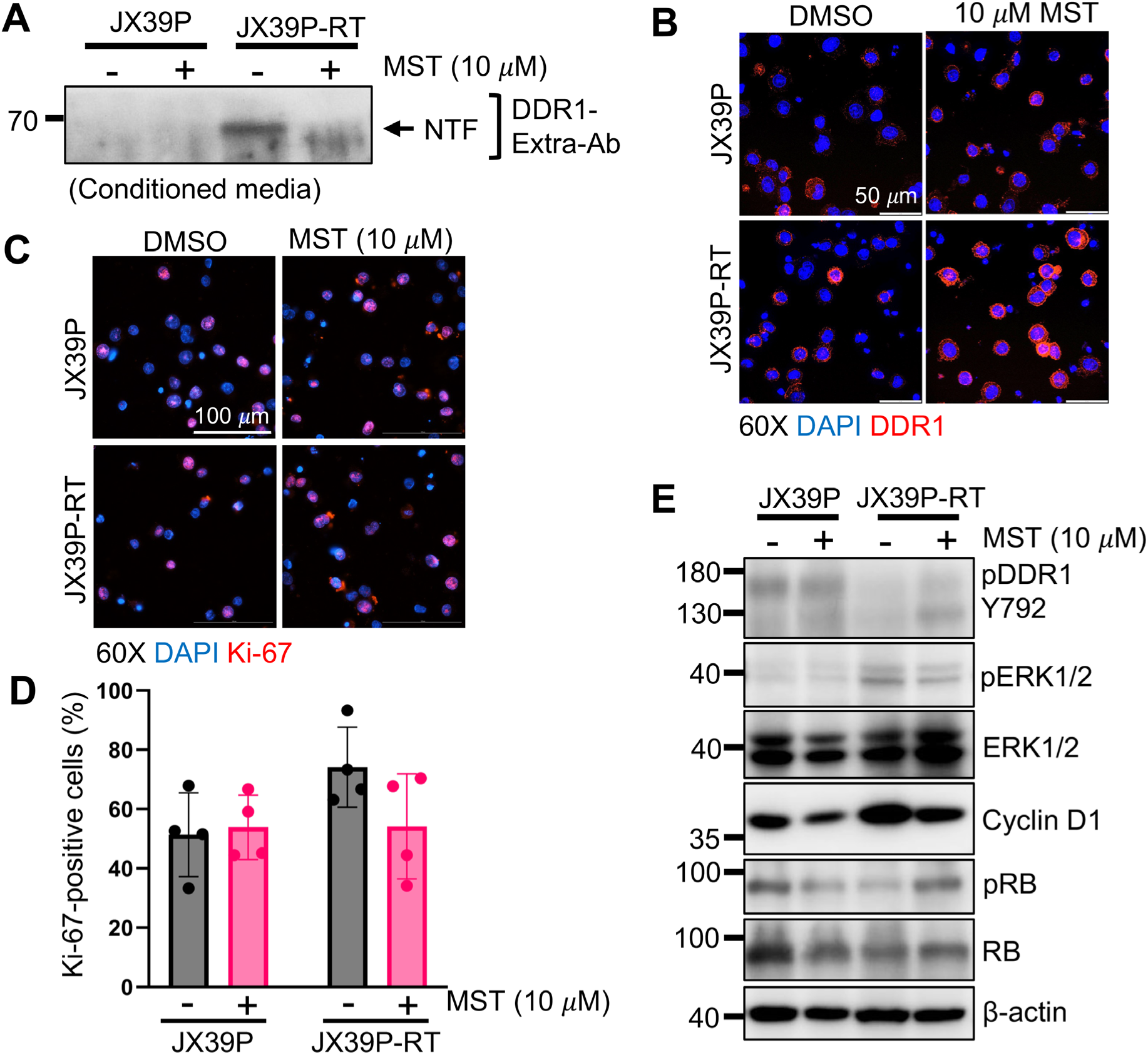
Broad metalloprotease inhibition suppresses DDR1 ectodomain shedding without enhancing proliferative signaling in radiation-resistant GBM. **(A)** Western blot analysis of concentrated conditioned media from collagen I-embedded JX39P and JX39P-RT cells treated with vehicle or the broad-spectrum metalloprotease inhibitor Marimastat (MST, 10 µM). MST treatment reduced the levels of shed DDR1 N-terminal fragment (NTF) in conditioned media from JX39P-RT cells. **(B)** Representative immunofluorescence images of DDR1 (red) and DAPI (blue) staining in JX39P and JX39P-RT cells following MST treatment, showing increased cell-associated DDR1 staining in JX39P-RT cells. Scale bars, 50 µm. **(C)** Representative immunofluorescence images of Ki-67 (red) and DAPI (blue) staining in JX39P and JX39P-RT cells treated with vehicle or MST. Scale bars, 100 µm. **(D)** Quantification of Ki-67-positive cells expressed as a percentage of total DAPI-positive nuclei. MST treatment did not significantly alter Ki-67 positivity in either cell line. **(E)** Western blot analysis of DDR1 phosphorylation at Y792, ERK1/2 phosphorylation, cyclin D1 expression, and RB phosphorylation in JX39P and JX39P-RT cells following MST treatment. MST increased DDR1 Y792 phosphorylation and RB phosphorylation without corresponding increases in ERK1/2 phosphorylation or cyclin D1 expression.

DDR1 IF staining further showed increased cell surface localization of DDR1 in MST-treated JX39P-RT cells, consistent with retention of cell-associated DDR1 following inhibition of ectodomain shedding (**Figure 4B**). However, Ki-67 staining revealed no significant change in the percentage of Ki-67-positive cells in either radiation-sensitive or radiation-resistant GBM following MST treatment, indicating that broad inhibition of metalloprotease activity does not enhance the proliferative activity of radiation-resistant cells (**Figure 4C, D**).

Western blot analysis showed increased DDR1 phosphorylation at Y792 following MST treatment, consistent with increased phosphorylation of cell-associated DDR1 when ectodomain shedding was suppressed. However, the increase in DDR1 phosphorylation was not accompanied by corresponding increases in ERK1/2 phosphorylation or cyclin D1 expression (**Figure 4E**). Although RB phosphorylation was increased following MST treatment, the absence of increased ERK1/2 phosphorylation, cyclin D1 expression, or Ki-67 positivity indicates that broad metalloprotease inhibition does not produce a coordinated increase in proliferative signaling.

Together, these findings demonstrate that pharmacologic inhibition of MMP activity reduces DDR1 ectodomain shedding and is associated with increased DDR1 phosphorylation in radiation-resistant GBM cells within a collagen I environment. However, these changes are not accompanied by a corresponding increase in ERK1/2 signaling or proliferative activity. Because MST inhibits multiple metalloproteases, these downstream effects cannot be attributed specifically to inhibition of DDR1 ectodomain shedding. These findings therefore support a role for metalloprotease activity in DDR1 shedding and underscore the need for a more specific approach to define the effects of DDR1 shedding on downstream signaling and proliferation in radiation-resistant GBM cells.

### *ADAMTS1* knockdown suppresses DDR1 ectodomain shedding and enhances proliferation signaling in radiation-resistant GBM

To identify candidate proteases associated with radiation resistance, we analyzed deep RNA sequencing data for JX39P/RT radiation-sensitive and radiation-resistant GBM PDX (40). Analysis of differentially expressed genes (DEGs) identified several MMP and ADAM family members that were upregulated in radiation-resistant GBM (**Figure 5A**). Among these candidates, *ADAMTS1* showed the strongest upregulation in radiation-resistant JX39P-RT relative to JX39P cells (adjusted p-value = 1.21 x 10-^6^), and this increase was confirmed by qRT-PCR (**Figure 5A, B**).

**Figure 5.**
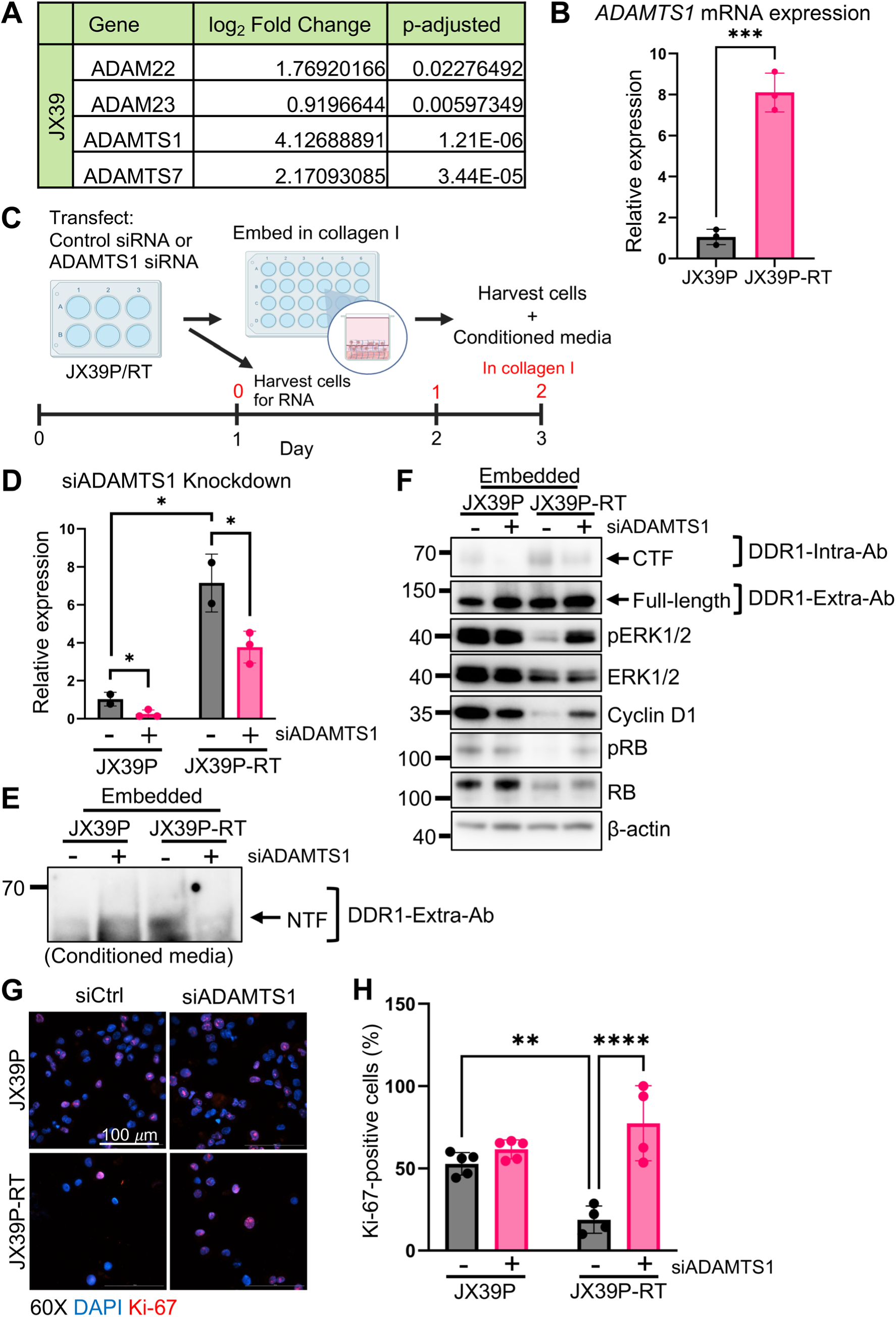
ADAMTS1 knockdown suppresses DDR1 ectodomain shedding and enhances proliferative signaling in radiation-resistant GBM. **(A)** Deep RNA sequencing analysis for MMP and ADAM family proteins upregulated in JX39P-RT. ADAMTS1 showed the greatest upregulation with log2 fold change = 4.13 and *p*-adjusted = 1.21x10^-6^. **(B)** Quantitative RT-PCR analysis of *ADAMTS1* mRNA expression in radiation-sensitive JX39P and radiation-resistant JX39P-RT cells, confirming increased *ADAMTS1* expression in JX39P- RT cells, consistent with RNA-sequencing data. \*\*\**p*<0.001 **(C)** Schematic illustrating the experimental timeline for siRNA-mediated ADAMTS1 knockdown and subsequent collagen I embedding. JX39P and JX39P-RT cells were transfected with control siRNA (siCtrl) or ADAMTS1-targeting siRNA (siADAMTS1) before collagen I embedding. Cell lysates and conditioned media were collected 72 h after transfection, corresponding to 2 days after collagen I embedding. **(D)** Quantitative RT-PCR analysis confirming *ADAMTS1* knockdown in JX39P and JX39P-RT cells before collagen I embedding. \**p*<0.05. **(E)** Western blot analysis of concentrated conditioned media from collagen I-embedded JX39P and JX39P-RT cells using an antibody recognizing the DDR1 extracellular domain (DDR1-Extra-Ab). ADAMTS1 knockdown reduced the levels of shed DDR1 N-terminal fragment (NTF) in conditioned media from JX39P-RT cells. **(F)** Western blot analysis of cell lysates from collagen I-embedded JX39P and JX39P-RT cells following ADAMTS1 knockdown. ADAMTS1 knockdown decreased generation of the DDR1 C-terminal fragment (CTF) and increased detection of full-length DDR1 in JX39P-RT cells, consistent with reduced DDR1 ectodomain shedding. Increased ERK1/2 and RB phosphorylation and cyclin D1 expression were also observed following ADAMTS1 knockdown in JX39P-RT cells. **(G)** Representative immunofluorescence images of Ki-67 (red) and DAPI (blue) staining in collagen I-embedded JX39P and JX39P-RT cells following transfection with siCtrl or siADAMTS1. Scale bars, 100 µm. **(H)** Quantification of Ki-67-positive cells expressed as a percentage of total DAPI-positive nuclei. ADAMTS1 knockdown increased the proportion of Ki-67-positive JX39P-RT cells compared with siCtrl-treated cells. \*\**p*<0.01, \*\*\*\**p*<0.0001.

To determine whether ADAMTS1 contributes to DDR1 ectodomain shedding, JX39P/RT cells were transfected with siRNA targeting ADAMTS1 (siADAMTS1) or control siRNA (siCtrl) and subsequently embedded in collagen I. After 2 days in the collagen I environment, cell lysates and conditioned media were collected for analysis (**Figure 5C**). To confirm siADAMTS1-mediated knockdown, cells were collected prior to embedding into collagen I. Downregulation of ADAMTS1 by siADAMTS1 was confirmed by qRT-PCR (**Figure 5D**). Consistent with our RNA-sequencing results, qRT-PCR also confirmed increased ADAMTS1 expression in JX39P-RT compared with JX39P cells (**Figure 5D**).

To assess DDR1 ectodomain shedding, conditioned media and cell lysates were analyzed for the shed DDR1 NTF and membrane-associated CTF, respectively. ADAMTS1 knockdown reduced the shed DDR1 NTF in conditioned media from JX39P-RT cells and decreased generation of the DDR1 CTF in cell lysates (**Figure 5E, F**). Concomitantly, analysis using an antibody recognizing the DDR1 extracellular domain showed increased detection of full-length DDR1 following ADAMTS1 knockdown (**Figure 5F**). Together, the reduced extracellular NTF, decreased CTF generation, and increased full-length DDR1 provide convergent evidence that ADAMTS1 contributes to DDR1 ectodomain shedding in radiation-resistant GBM cells.

We next examined whether ADAMTS1 knockdown was accompanied by changes in proliferative signaling. In JX39P-RT cells, ADAMTS1 knockdown increased ERK1/2 and RB phosphorylation as well as cyclin D1 expression (**Figure 5F**). Consistent with these signaling changes, Ki-67 staining revealed an increased percentage of Ki-67-positive JX39P-RT cells following ADAMTS1 knockdown (**Figure 5G, H**). Therefore, ADAMTS1 knockdown simultaneously reduced DDR1 ectodomain shedding and enhanced proliferative signaling and activity in radiation-resistant GBM cells.

Together, these data identify ADAMTS1 as a regulator contributing to DDR1 ectodomain shedding in radiation-resistant GBM. ADAMTS1 knockdown reduces both release of the DDR1 NTF and generation of the membrane-associated CTF while increasing full-length DDR1, and these changes are accompanied by increased ERK1/2 and RB phosphorylation, cyclin D1 expression, and Ki-67 positivity in JX39P-RT cells. Collectively, these findings support a proposed model in which enhanced ADAMTS1-associated DDR1 ectodomain shedding is linked to attenuated proliferative signaling in acquired radiation-resistant GBM cells cultured in a 3D collagen I matrix (**Figure 6**).

**Figure 6.**
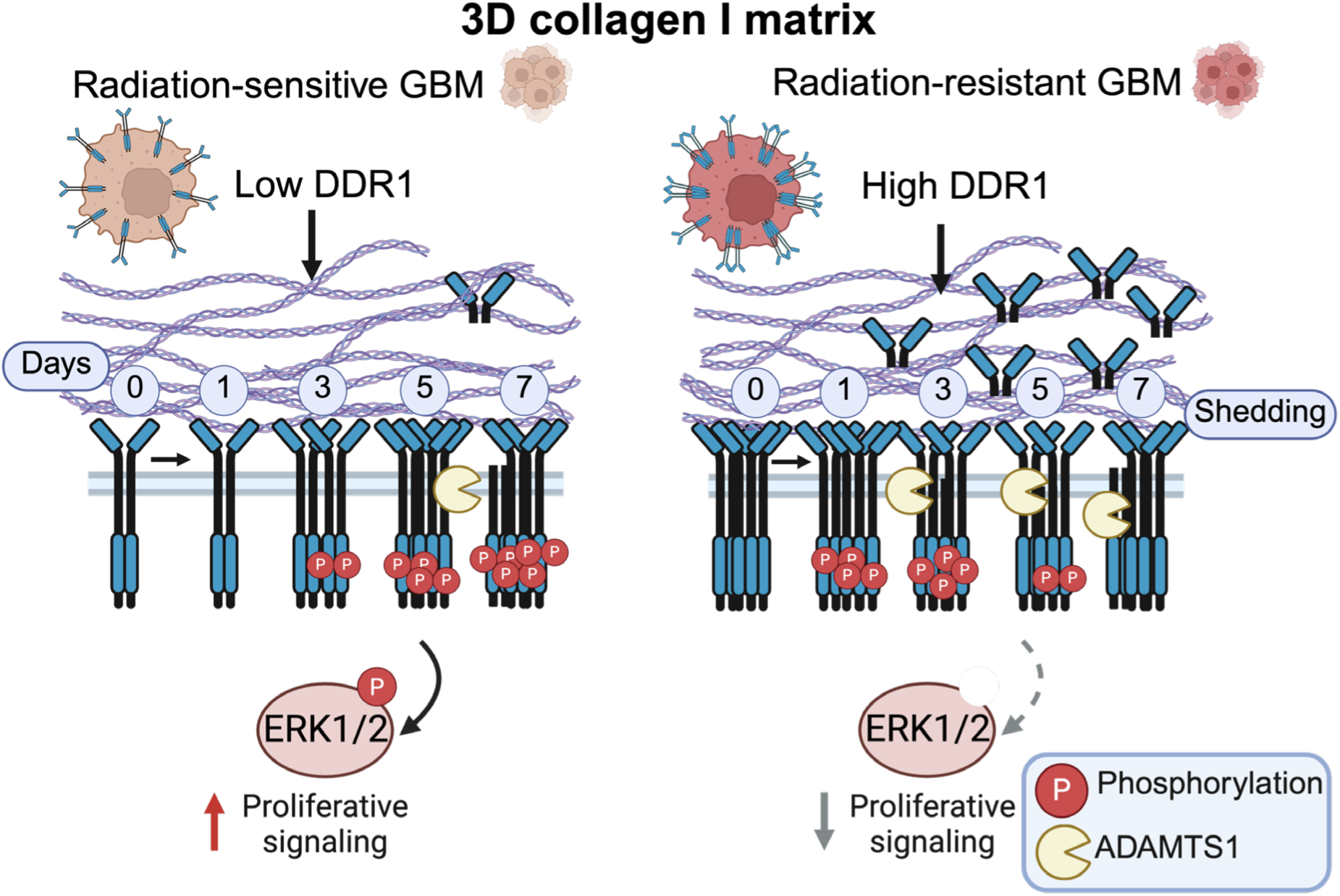
Proposed model of enhanced DDR1 ectodomain shedding and attenuated proliferative signaling in radiation-resistant GBM. Schematic illustrating distinct DDR1 processing and signaling states in radiation-sensitive parental and acquired radiation-resistant GBM cells cultured in a three-dimensional (3D) collagen I matrix. Compared with parental cells, radiation-resistant GBM cells exhibit increased DDR1 expression, enhanced ectodomain shedding, and reduced receptor phosphorylation. Enhanced shedding is characterized by increased release of the DDR1 N-terminal fragment (NTF) and accumulation of the membrane-associated C-terminal fragment (CTF). ADAMTS1 is upregulated in radiation-resistant cells, and its knockdown reduces DDR1 shedding while increasing full-length DDR1 abundance and ERK1/2 phosphorylation, accompanied by enhanced cell cycle-associated signaling and Ki-67 positivity. The model proposes an association between enhanced ADAMTS1-associated DDR1 shedding and attenuated proliferative signaling in radiation-resistant GBM. Dashed arrows indicate proposed signaling relationships that remain to be mechanistically established.

## DISCUSSION

Radiation resistance remains a barrier to effectively treating GBM and is a major contributor to tumor recurrence and poor patient prognosis (2–4). The development of effective treatments is complicated by inherent features of GBM, including tumor heterogeneity, the blood-brain barrier, and an immunosuppressive tumor microenvironment (8–11). While tumor-intrinsic mechanisms underlying radiation resistance have been extensively studied, the contribution of the tumor microenvironment remains poorly understood. ECM remodeling is an important component of the tumor microenvironment that can influence therapeutic response. Although DDR1 ectodomain shedding and its influence on receptor signaling have been reported in other cellular contexts, how DDR1 processing differs between radiation-sensitive and acquired radiation-resistant GBM remains unclear. Here, we identify a distinct DDR1 regulatory state in radiation-resistant GBM cells embedded in a 3D collagen I matrix, characterized by enhanced receptor shedding despite increased DDR1 expression, reduced DDR1 phosphorylation, and altered proliferative signaling.

Our analysis revealed elevated COL1A1 and COL1A2 mRNA levels and COL1A2 protein abundance in GBM compared with normal brain tissue. When embedded in a 3D collagen I matrix, radiation-resistant GBM cells exhibited a distinct temporal pattern of Ki-67 positivity, with higher levels at early time points followed by a decline during prolonged culture. Radiation-sensitive and radiation-resistant GBM cells also exhibited distinct temporal patterns of ERK1/2 phosphorylation following collagen I embedding, indicating differential signaling responses under these culture conditions. Together, these findings demonstrate that proliferative activity and ERK1/2 signaling differ between radiation-sensitive and radiation-resistant GBM cells within a defined 3D collagen I matrix, highlighting the potential importance of ECM context in shaping the behavior of radiation-resistant GBM.

We demonstrate that DDR1 is upregulated at both the mRNA and protein levels in radiation-resistant GBM. This dysregulation underscores the molecular differences between radiation-sensitive and radiation-resistant GBM and the importance of post-translational regulation, as reflected in DDR1’s regulation through ligand bioavailability, proteolytic processing, and receptor-membrane dynamics (29–33). As a collagen-activated receptor tyrosine kinase, DDR1 provides a potential mechanism through which the collagen microenvironment influences intracellular signaling. Notably, despite increased DDR1 expression in radiation-resistant JX39P-RT cells, DDR1 phosphorylation at Y792 and Y513 was markedly lower than in parental JX39P cells following collagen I embedding, indicating that increased receptor abundance does not necessarily result in increased DDR1 activation. DDR1 KO experiments further demonstrated that loss of DDR1 reduces ERK1/2 phosphorylation, supporting a role for DDR1 in ERK1/2 signaling within the 3D collagen I matrix.

The central finding of this study is that acquired radiation-resistant GBM cells exhibit enhanced DDR1 ectodomain shedding despite increased DDR1 expression, accompanied by markedly reduced receptor phosphorylation within a 3D collagen I matrix. Although DDR1 ectodomain shedding has previously been described in other cellular systems, our findings reveal that DDR1 processing is distinctly altered in radiation-resistant GBM compared with matched parental cells. Increased accumulation of the membrane-associated C-terminal fragment and release of the N-terminal fragment indicate enhanced receptor processing in the resistant state. These observations suggest that increased DDR1 expression does not necessarily translate into increased receptor activation and raise the possibility that enhanced ectodomain shedding contributes to reduced availability of intact DDR1 for collagen-dependent signaling. Previous studies implicating the shed DDR1 ectodomain in collagen organization and immune exclusion further suggest that enhanced DDR1 shedding may have biological consequences beyond receptor attenuation (34, 35). Thus, the potential extracellular functions of the shed DDR1 NTF in radiation-resistant GBM warrant further investigation.

To investigate whether pharmacologic suppression of DDR1 ectodomain shedding alters DDR1 signaling and proliferative activity, we used MST, a broad-spectrum metalloprotease inhibitor. MST reduced DDR1 ectodomain shedding and increased cell-associated DDR1 and DDR1 phosphorylation. However, increased DDR1 phosphorylation was not accompanied by corresponding increases in ERK1/2 phosphorylation, cyclin D1 expression, or Ki-67 positivity, despite increased RB phosphorylation. This discordant response suggests that broad metalloprotease inhibition affects multiple signaling pathways and limits interpretation of MST treatment as a specific test of DDR1 ectodomain shedding. While broad metalloprotease inhibitors have not had great success in the clinic (41),these findings underscore the importance of identifying specific proteases that regulate DDR1 shedding to more directly define its functional consequences.

To identify a more specific regulator of DDR1 ectodomain shedding, we analyzed RNA sequencing data from matched radiation-sensitive and radiation-resistant GBM PDX models. RNA sequencing revealed broad upregulation of metalloprotease and ADAM family genes, with ADAMTS1 showing the most significant increase in JX39P-RT cells. *ADAMTS1* knockdown reduced NTF release into conditioned media and CTF generation in cell lysates while increasing cell-associated full-length DDR1, providing convergent evidence that ADAMTS1 contributes to DDR1 ectodomain shedding in radiation-resistant GBM. Following *ADAMTS1* knockdown, increased phosphorylation of ERK1/2 and RB and cyclin D1 expression was detected, along with increased Ki-67 positivity in radiation-resistant GBM. These data identify ADAMTS1 as a regulator of DDR1 proteolytic processing in radiation-resistant GBM and suggest that additional metalloproteases may also participate in this process. This aligns with previous studies implicating metalloprotease-mediated ECM remodeling in aggressive tumor phenotypes and provides a basis for further investigation of protease-dependent ECM signaling in radiation-resistant GBM (18, 19). Importantly, the concomitant reduction in DDR1 ectodomain shedding and increase in proliferative signaling following ADAMTS1 knockdown support a model in which enhanced DDR1 shedding is associated with attenuation of proliferative signaling in radiation-resistant GBM cells. Whether this shift in proliferative state alters radiation sensitivity remains to be determined.

A limitation of this study is that although the 3D collagen I matrix provides a defined experimental model for investigating collagen-dependent DDR1 regulation, it does not fully replicate the complexity of the brain ECM, which also contains hyaluronan, laminins, and proteoglycans. Future work should incorporate more complex ECM models to determine how additional ECM components influence DDR1 ectodomain shedding and signaling. In addition, we identified ADAMTS1 as a regulator that contributes to DDR1 ectodomain shedding; however, other proteases may also cleave DDR1 and require further investigation. Furthermore, although *ADAMTS1* knockdown reduced DDR1 ectodomain shedding and was accompanied by increased proliferative signaling, the present study does not establish that these signaling changes are mediated specifically through DDR1. Future studies combining ADAMTS1 depletion with DDR1 loss or inhibition will be important to determine the DDR1 dependence of this phenotype.

In summary, our findings, illustrated in **Figure 6**, identify enhanced DDR1 ectodomain shedding as a prominent feature of acquired radiation-resistant GBM cells cultured in a 3D collagen I matrix, associated with altered receptor activation and proliferative signaling. Despite increased DDR1 expression, radiation-resistant cells exhibit enhanced receptor shedding and reduced DDR1 phosphorylation, whereas ADAMTS1 knockdown reduces shedding and is accompanied by increased full-length DDR1 abundance and enhanced ERK1/2 and cell cycle-associated signaling. Together, these findings support a model in which enhanced DDR1 ectodomain shedding is linked to attenuation of proliferative signaling in radiation-resistant GBM. This study identifies altered collagen-dependent DDR1 processing as a previously unexplored post-translational regulatory feature of acquired radiation-resistant GBM and provides a basis for investigating whether modulation of DDR1 shedding influences therapeutic response.

## MATERIALS AND METHODS

### Cell culture

The PDX line, GBM39 (designated JX39P), was originally obtained from JN Sarkaria (Mayo Clinic, Rochester, Minnesota, USA) and maintained by the UAB Brain Tumor Animal Model Core. JX39P (radiation-sensitive) and JX39P-RT (radiation-resistant) GBM PDX models were maintained and generated, respectively, in the laboratory of Dr. Christopher Willey. The radiation-resistant JX39P-RT was generated by serial *in vivo* irradiation. Mice received 6 fractions of 2 Gy until the tumor doubling time was less than 20 days, as described in more detail (36). HEK293T cells, a human embryonic kidney-derived cell line, were used for viral production.

All cell cultures were maintained at 37℃ in 5% CO_2_. JX39P and JX39P-RT GBM PDX cells were maintained in liquid suspension and grown in GBM stem cell media consisting of DMEM/F12 (Elabscience #PM150310) supplemented with Gem21 Neuroplex (GeminiBio #400-161-010), 20 ng/mL human EGF (SinoBiological 10605-HNAE), 20 ng/mL human bFGF (SinoBiological 10014-HNAE), and 5 µg/mL heparin (MedChemExpress HY-17567C). HEK293T cells were grown in DMEM (Elabscience #PM150210) supplemented with 10% FBS (GeminiBio #S11150).

### Generation of DDR1 Knockout GBM PDX Cells

HEK293T cells were seeded at 60% confluency. Second-generation lentiviral packaging vectors, psPAX2 (13.5 *μ*g) and pMD2.G (6.5 *μ*g), were co-transfected with 20 *μ*g of Tet-inducible Cas9 lentiviral vector, pCW-Cas9-Blast (Addgene #83481). After 8 hours, the media was changed from DMEM with 10% FBS to GBM stem cell media supplemented with Gem21 only. After 24 hours, the virus was harvested, and 10 mL was added to each plate of JX39P/RT cells. JX39P/RT cells were selected with 10 µg/mL blasticidin (Goldbio #B-800-100).

We repeated the infection protocol in JX39P/RT-Cas9 cells using either lentiGuide-Puro (Addgene #52963) or DDR1 gRNA (Addgene #78012). Cells were selected with 10 µg/mL blasticidin and 1 µg/mL puromycin (Goldbio #P-600-100). Cells were treated with 1 µg/mL doxycycline (Goldbio #D-500-1) to induce Cas9 expression and promote DDR1 knockout (KO). Western blot confirmed DDR1 KO.

### Analysis of DDR1 Expression in TCGA and CPTAC Datasets

DDR1 protein and mRNA expression data were obtained and visualized using the **U**niversity of **AL**abama at Birmingham **CAN**cer data analysis Portal (UALCAN) (42, 43). In UALCAN, the GBM dataset was selected for TCGA data analysis, and GBM (Extended) was selected for CPTAC data analysis.

### RNA isolation and cDNA synthesis

RNA was isolated using TRIzol (Thermo Fisher Scientific #15596018) following the standard protocol. RNA was DNase 1 treated using the DNA-free Kit (Thermo Fisher Scientific #AM1906). cDNA was synthesized using the SuperScript III First-Strand Synthesis System for RT-PCR (Thermo Fisher Scientific #18080051), following the standard product protocol. However, 0.5 *μ*L per sample of SuperScript III and RNase OUT enzymes were used.

### Quantitative PCR and analysis

Quantitative PCR (qRT-PCR) was performed on a Bio-Rad CFX Connect machine using the following protocol: denaturation at 95℃ for 3 minutes; 40 cycles of denaturation at 95℃ for 10 seconds and annealing/elongation at 60℃ for 30 seconds; and a melting curve at 65℃ for 5 seconds and 95℃ for 50 seconds. iTaq SYBR (Bio-Rad #1725124) was mixed with primers to make a master mix, and cDNA was added separately. Samples were tested in triplicate. Quantification was performed in Excel and normalized to the Control siRNA (siCtrl) or radiation-sensitive sample. Relative expression (2^-ΔΔCt^) is graphed using Prism.

Primers were designed with the IDT PrimerQuest tool. *DDR1* primers have the forward primer in coding exon 1 and the reverse primer in coding exon 2. Primers are listed 5’ to 3’: DDR1_F: TGGAGATGCTGACATGAAGG, DDR1_R: GAGATGTCACTGTCTGGGATG. The *ADAMTS1*forward and reverse primers are in coding exons 8 and 9, respectively. Primers are listed 5’ to 3’: ADAMTS1_F: ATCAGGATCAGTTACTAGTGCAAA, ADAMTS1_R: CTCTGGTTCCGCTGTTTCA. The TUBA1A primers are on coding exon 2 (forward) and 3 (reverse). TUBA1A_F: GGCAGTGTTTGTAGACTTGGA, TUBA1A_R: GCATCTTCTTTGCCTGTGATAAG.

### Western blot analyses

SDS-PAGE gels were made following standard recipes. To detect DDR1, pRB, and RB, 8% resolving gels were used. β-actin, pERK1/2, ERK1/2, and cyclin D1 were detected from 12% resolving gels. All gels were run at 100V. Proteins of interest run on 8% gels were transferred for 1 hour and 45 minutes at 100 V, and 12% gels were transferred for 1 hour.

Primary antibodies were diluted in 5% BSA and incubated overnight at 4℃. Antibodies were diluted as follows: DDR1-Intra-Ab (Cell Signaling #5583S) 1:1,000, DDR1 phospho-Y513 (Cell Signaling #14531S) 1:1,000, DDR1 phospho-Y792 (Cell Signaling #11994S) 1:1,000, DDR1- Extra-Ab (Santa Cruz #sc-390268) 1:500, β-actin (catalog) 1:10,000, pERK1/2 (Cell Signaling #9101S) 1:2,000, ERK1/2 (Cell Signaling #4695S) 1:5,000, pRB (Protein Tech #30376-1-AP) 1:1,000, RB (Protein Tech #10048-2-Ig) 1:1,000, cyclin D1 (Thermo Fisher Scientific #MA5-14512) 1:1,000.

Secondary antibodies were diluted in 5% BSA and incubated at room temperature for 1 hour and 30 minutes. Rabbit secondary (Protein Tech #SA00001-2) was diluted to 1:10,000, and mouse secondary (Thermo Fisher Scientific #31430) was diluted to 1:10,000.

### Immunofluorescence

Cells were suspended in HBSS (Cytiva #SH30268.02) at 100K cells in 1 mL. The Cytospin 3 machine (Shandon) was used to deposit the cells onto the slide. Slides were spun at 800 rpm for 3 minutes and fixed in methanol.

Slides were rehydrated in 1X PBS for 3 × 10 minutes on a rocker and placed in blocking buffer (2% BSA, 0.1% Triton in PBS) on the rocker for 45 minutes. Slides were incubated in primary antibody for 1 hour. Primary antibodies were diluted in blocking buffer: DDR1 (Cell Signaling #5583S) 1:200, Ki-67 (DSHB #AFFN-Ki67-3E6) 1:200. Slides were washed in 0.2% TBST for 3 × 10 minutes on a rocker. Secondary antibody and DAPI were diluted in blocking buffer: Alexa 594-Rabbit (Thermo Fisher Scientific #A11012) 1:1,000, Alexa 594-Mouse (Thermo Fisher Scientific #A11005) 1:500, and DAPI (Therm Fisher Scientific #62248) 1:1,000. Secondary antibody with DAPI was incubated for 30 minutes. Slides were washed again in 0.2% PBST for 3 × 10 minutes. DDR1 IF images were taken on the CrestOptics X-Light V3 Spinning Disk Confocal Imager with an Olympus Ix83 lens at 60X with oil, and representative images are presented here. Ki-67 IF images were taken on Agilent BioTek Lionheart LX Imager at 60X with oil.

Ki-67 quantification was performed using the Gen5 program on the Lionheart Imager. The percentage of the Ki-67-positive subpopulation was determined based on total nuclei counted in at least 3 fields of view. A threshold of 5000 dpi was used to indicate positive Ki-67 staining.

### Collagen I-embedded culture and harvest from collagen matrix

Collagen base was prepared on ice with 5.5 mL collagen type I (Gibco #A10483-01), 100 mM HEPES (Corning #25-060-CI), and 275 µL GBM stem cell media. 200 µL of this mixture was added to each well in a 24-well plate and allowed to solidify in the cell culture incubator at 37℃.

JX39P/RT cells were counted with a Countess 3 FL to embed the same number of cells in each plate and suspended in 8.166 mL GBM stem cell media.

After the first base layer solidified, the second collagen I layer was prepared with 7 mL collagen I, 100 mM HEPES, and 350 µL GBM stem cell media. Cells from before were combined with the second collagen layer mix, and 500 *μ*L was added to each well. Plates were allowed to solidify for up to 1 hour before adding GBM stem cell media.

To recover cells from the collagen matrix, 0.3 mg/mL collagenase type I (Gibco #17100-017) was prepared fresh in room-temperature HBSS. Collagen disks were washed with HBSS and put in 1 mL of collagenase each. The collagen disks were incubated in collagenase for ∼45 minutes on a rocker at 37℃. After recovery, cells were collected for further experiments: IF slides or protein lysates. Conditioned media were collected to detect the DDR1 ectodomain fragment.

### Collagen I time-course experiments

For time-course experiments, JX39P and JX39P-RT cells were embedded in collagen I as described above and harvested 1, 3, 5, and 7 days after embedding. Cells maintained in liquid suspension culture without collagen I were analyzed as controls.

### DDR1-IN-1 treatment

For inhibition of DDR1 kinase activity, DDR1-IN-1 (10 µM; MedChemExpress, HY-13979) was included in both the cell-containing second collagen layer and the overlying culture medium. Cells were harvested 24 h after collagen I embedding for subsequent analysis.

### Marimastat treatment

For broad metalloprotease inhibition, Marimastat (10 µM; MedChemExpress, HY-12169) was included in both the cell-containing second collagen layer and the overlying culture medium. Cells and conditioned medium were collected 3 days after collagen I embedding for subsequent analyses.

### Collagen I embedded and ADAMTS1 siRNA experiment

JX39P/RT cells were reverse-transfected with siRNA, and 5 wells (6-well plate) per condition were prepared: JX39P siCtrl, JX39P siADAMTS1, JX39P-RT siCtrl, JX39P-RT siADAMTS1. JX39P/RT cells were counted, and 400K cells per well were resuspended to add 1.75 mL to each well.

Following guidelines for RNAiMAX (Thermo Fisher Scientific #13778150), JX39P/RT cells were treated with either 80 pmol of Control siRNA or *ADAMTS1* siRNA. Control siRNA is from Thermo Fisher (Thermo Fisher #4390844), and *ADAMTS1* siRNA is from IDT (IDT #574921982), and the duplex sequence is 5’ AGUGGUCUAAAGCAUUACCUUCUCA 3’ and 5’ UGAGAAGGUAAUGCUUUAGACCACUGC 3’. Cells were incubated overnight.

Cells were prepared for embedding as described above or reserved for RNA isolation. Cells were harvested from the collagen I matrix on day 2 (72 hours after siRNA treatment) following the above protocol; samples were collected for protein lysates, IF slides, and conditioned media. qRT-PCR confirmed *ADAMTS1* knockdown.

### Conditioned media concentration and sample preparation

From the collected conditioned media (same volume), 20 mL of each sample was added to protein concentrators (Pierce #88526) and centrifuged at 4,000 rpm until concentrated to 200 µL. 2XSDS Laemmli buffer was added proportionally.

### Differentially Expressed Gene (DEG) analysis of paired GBM PDXs

Deep RNA sequencing for paired GBM PDX JX39P/RT was analyzed for *MMP* and *ADAM* genes. Genes were filtered for positive log2 fold change and were considered significant with p-adjusted < 0.05.

### Statistical analysis

Experiments were repeated in triplicate, and all statistical analyses, including t-tests and 2-way ANOVAs, were performed in Prism. P-values < 0.05 were considered significant.

## ACKNOWLEDGEMENTS

This work was supported by the National Institutes of Health (NIH) (R01CA236911 and R01HL168659 to E.E.A.; R01HL158800 and R01HL158875 to S.S.L.; U01CA223976 and 3U01CA223976-03S1 to C.D.W.), the UAB Bridge Fund (to C.D.W.), the University of Alabama at Birmingham (UAB) Heersink School of Medicine, UAB Department of Pathology, and UAB O’Neal Cancer Center’s O’Neal Invests Program (Catalyst grant to C.D.W. and E.E.A.). S.D., C.A.H., and T.L.S. were supported by the NIH Training Program T32 (grant no. T32GM135028 for S.D.; grant no. T32CA047888 for C.A.H.; grant no. T32NS121721 for T.L.S.). This content is solely the responsibility of the authors and does not necessarily represent the views of the NIH.

## REFERENCES

1. Melhem JM, Detsky J, Lim-Fat MJ, Perry JR. Updates in IDH-Wildtype Glioblastoma. Neurotherapeutics. 2022;19(6):1705–23.

2. Wen PY, Weller M, Lee EQ, Alexander BM, Barnholtz-Sloan JS, Barthel FP, Batchelor TT, Bindra RS, Chang SM, Chiocca EA, Cloughesy TF, DeGroot JF, Galanis E, Gilbert MR, Hegi ME, Horbinski C, Huang RY, Lassman AB, Le Rhun E, Lim M, Mehta MP, Mellinghoff IK, Minniti G, Nathanson D, Platten M, Preusser M, Roth P, Sanson M, Schiff D, Short SC, Taphoorn MJB, Tonn JC, Tsang J, Verhaak RGW, von Deimling A, Wick W, Zadeh G, Reardon DA, Aldape KD, van den Bent MJ. Glioblastoma in adults: a Society for Neuro-Oncology (SNO) and European Society of Neuro-Oncology (EANO) consensus review on current management and future directions. Neuro Oncol. 2020;22(8):1073–113.

3. Sipos D, Raposa BL, Freihat O, Simon M, Mekis N, Cornacchione P, Kovacs A. Glioblastoma: Clinical Presentation, Multidisciplinary Management, and Long-Term Outcomes. Cancers (Basel). 2025;17(1).

4. Birzu C, French P, Caccese M, Cerretti G, Idbaih A, Zagonel V, Lombardi G. Recurrent Glioblastoma: From Molecular Landscape to New Treatment Perspectives. Cancers (Basel). 2020;13(1).

5. Aldape K, Brindle KM, Chesler L, Chopra R, Gajjar A, Gilbert MR, Gottardo N, Gutmann DH, Hargrave D, Holland EC, Jones DTW, Joyce JA, Kearns P, Kieran MW, Mellinghoff IK, Merchant M, Pfister SM, Pollard SM, Ramaswamy V, Rich JN, Robinson GW, Rowitch DH, Sampson JH, Taylor MD, Workman P, Gilbertson RJ. Challenges to curing primary brain tumours. Nat Rev Clin Oncol. 2019;16(8):509–20.

6. Hegi ME, Diserens AC, Gorlia T, Hamou MF, de Tribolet N, Weller M, Kros JM, Hainfellner JA, Mason W, Mariani L, Bromberg JE, Hau P, Mirimanoff RO, Cairncross JG, Janzer RC, Stupp R. MGMT gene silencing and benefit from temozolomide in glioblastoma. N Engl J Med. 2005;352(10):997–1003.

7. Bonm A, Kesari S. DNA Damage Response in Glioblastoma: Mechanism for Treatment Resistance and Emerging Therapeutic Strategies. Cancer J. 2021;27(5):379–85.

8. Eisenbarth D, Wang YA. Glioblastoma heterogeneity at single cell resolution. Oncogene. 2023;42(27):2155–65.

9. Moreno-Sanchez PM, Rezaeipour M, Inderberg EM, Platten M, Golebiewska A. Immunosuppressive mechanisms and therapeutic interventions shaping glioblastoma immunity. Nat Cancer. 2026;7(1):29–42.

10. Arvanitis CD, Ferraro GB, Jain RK. The blood-brain barrier and blood-tumour barrier in brain tumours and metastases. Nat Rev Cancer. 2020;20(1):26–41.

11. Goenka A, Tiek D, Song X, Huang T, Hu B, Cheng SY. The Many Facets of Therapy Resistance and Tumor Recurrence in Glioblastoma. Cells. 2021;10(3).

12. Karamanos NK, Theocharis AD, Piperigkou Z, Manou D, Passi A, Skandalis SS, Vynios DH, Orian-Rousseau V, Ricard-Blum S, Schmelzer CEH, Duca L, Durbeej M, Afratis NA, Troeberg L, Franchi M, Masola V, Onisto M. A guide to the composition and functions of the extracellular matrix. FEBS J. 2021;288(24):6850–912.

13. Cordes N, Blaese MA, Plasswilm L, Rodemann HP, Van Beuningen D. Fibronectin and laminin increase resistance to ionizing radiation and the cytotoxic drug Ukrain in human tumour and normal cells in vitro. Int J Radiat Biol. 2003;79(9):709–20.

14. Pu X, Chen K, Dong L, Yi J, Wang M, Wei X, Zhao M, Zhao M, Wang X, Ding L, Jin S. FHL2 enhances ITGB1-mediated ECM remodeling and cellular stiffness to promote radioresistance in non-small cell lung cancer. Cell Death Discov. 2025;11(1):480.

15. Baker AM, Bird D, Lang G, Cox TR, Erler JT. Lysyl oxidase enzymatic function increases stiffness to drive colorectal cancer progression through FAK. Oncogene. 2013;32(14):1863–8.

16. Mao M, Li Z, Zheng W, Xia H, Fei Z, Yao Q, Zheng R. Collagen type I alpha 1 induces radio-resistance in hypoxic glioblastoma cells by promoting autophagy. J Neurooncol. 2025;175(1):287–301.

17. Chernov AV, Baranovskaya S, Golubkov VS, Wakeman DR, Snyder EY, Williams R, Strongin AY. Microarray-based transcriptional and epigenetic profiling of matrix metalloproteinases, collagens, and related genes in cancer. J Biol Chem. 2010;285(25):19647–59.

18. Niland S, Riscanevo AX, Eble JA. Matrix Metalloproteinases Shape the Tumor Microenvironment in Cancer Progression. Int J Mol Sci. 2021;23(1).

19. Mohiuddin E, Wakimoto H. Extracellular matrix in glioblastoma: opportunities for emerging therapeutic approaches. Am J Cancer Res. 2021;11(8):3742–54.

20. Berestjuk I, Lecacheur M, Carminati A, Diazzi S, Rovera C, Prod’homme V, Ohanna M, Popovic A, Mallavialle A, Larbret F, Pisano S, Audebert S, Passeron T, Gaggioli C, Girard CA, Deckert M, Tartare-Deckert S. Targeting Discoidin Domain Receptors DDR1 and DDR2 overcomes matrix-mediated tumor cell adaptation and tolerance to BRAF-targeted therapy in melanoma. EMBO Mol Med. 2022;14(2):e11814.

21. Shrivastava A, Radziejewski C, Campbell E, Kovac L, McGlynn M, Ryan TE, Davis S, Goldfarb MP, Glass DJ, Lemke G, Yancopoulos GD. An orphan receptor tyrosine kinase family whose members serve as nonintegrin collagen receptors. Mol Cell. 1997;1(1):25–34.

22. Vogel W, Gish GD, Alves F, Pawson T. The discoidin domain receptor tyrosine kinases are activated by collagen. Mol Cell. 1997;1(1):13–23.

23. Curat CA, Eck M, Dervillez X, Vogel WF. Mapping of epitopes in discoidin domain receptor 1 critical for collagen binding. J Biol Chem. 2001;276(49):45952–8.

24. Carafoli F, Hohenester E. Collagen recognition and transmembrane signalling by discoidin domain receptors. Biochim Biophys Acta. 2013;1834(10):2187–94.

25. Noordeen NA, Carafoli F, Hohenester E, Horton MA, Leitinger B. A transmembrane leucine zipper is required for activation of the dimeric receptor tyrosine kinase DDR1. J Biol Chem. 2006;281(32):22744–51.

26. Juskaite V, Corcoran DS, Leitinger B. Collagen induces activation of DDR1 through lateral dimer association and phosphorylation between dimers. Elife. 2017;6.

27. Corcoran DS, Juskaite V, Xu Y, Gorlitz F, Alexandrov Y, Dunsby C, French PMW, Leitinger B. DDR1 autophosphorylation is a result of aggregation into dense clusters. Sci Rep. 2019;9(1):17104.

28. Leitinger B. Discoidin domain receptor functions in physiological and pathological conditions. Int Rev Cell Mol Biol. 2014;310:39–87.

29. Fu HL, Valiathan RR, Arkwright R, Sohail A, Mihai C, Kumarasiri M, Mahasenan KV, Mobashery S, Huang P, Agarwal G, Fridman R. Discoidin domain receptors: unique receptor tyrosine kinases in collagen-mediated signaling. J Biol Chem. 2013;288(11):7430–7.

30. Mihai C, Chotani M, Elton TS, Agarwal G. Mapping of DDR1 distribution and oligomerization on the cell surface by FRET microscopy. J Mol Biol. 2009;385(2):432–45.

31. Shitomi Y, Thogersen IB, Ito N, Leitinger B, Enghild JJ, Itoh Y. ADAM10 controls collagen signaling and cell migration on collagen by shedding the ectodomain of discoidin domain receptor 1 (DDR1). Mol Biol Cell. 2015;26(4):659–73.

32. Slack BE, Siniaia MS, Blusztajn JK. Collagen type I selectively activates ectodomain shedding of the discoidin domain receptor 1: involvement of Src tyrosine kinase. J Cell Biochem. 2006;98(3):672–84.

33. Vogel WF. Ligand-induced shedding of discoidin domain receptor 1. FEBS Lett. 2002;514(2-3):175–80.

34. Sun X, Wu B, Chiang HC, Deng H, Zhang X, Xiong W, Liu J, Rozeboom AM, Harris BT, Blommaert E, Gomez A, Garcia RE, Zhou Y, Mitra P, Prevost M, Zhang D, Banik D, Isaacs C, Berry D, Lai C, Chaldekas K, Latham PS, Brantner CA, Popratiloff A, Jin VX, Zhang N, Hu Y, Pujana MA, Curiel TJ, An Z, Li R. Tumour DDR1 promotes collagen fibre alignment to instigate immune exclusion. Nature. 2021;599(7886):673–8.

35. Liu J, Chiang HC, Xiong W, Laurent V, Griffiths SC, Dulfer J, Deng H, Sun X, Yin YW, Li W, Audoly LP, An Z, Schurpf T, Li R, Zhang N. A highly selective humanized DDR1 mAb reverses immune exclusion by disrupting collagen fiber alignment in breast cancer. J Immunother Cancer. 2023;11(6).

36. Stackhouse CT, Anderson JC, Yue Z, Nguyen T, Eustace NJ, Langford CP, Wang J, Rowland JRt, Xing C, Mikhail FM, Cui X, Alrefai H, Bash RE, Lee KJ, Yang ES, Hjelmeland AB, Miller CR, Chen JY, Gillespie GY, Willey CD. An in vivo model of glioblastoma radiation resistance identifies long noncoding RNAs and targetable kinases. JCI Insight. 2022;7(16).

37. Dumesic PA, Scholl FA, Barragan DI, Khavari PA. Erk1/2 MAP kinases are required for epidermal G2/M progression. J Cell Biol. 2009;185(3):409–22.

38. Meloche S, Pouyssegur J. The ERK1/2 mitogen-activated protein kinase pathway as a master regulator of the G1-to S-phase transition. Oncogene. 2007;26(22):3227–39.

39. Baltes F, Caspers J, Henze S, Schlesinger M, Bendas G. Targeting Discoidin Domain Receptor 1 (DDR1) Signaling and Its Crosstalk with beta(1)-integrin Emerges as a Key Factor for Breast Cancer Chemosensitization upon Collagen Type 1 Binding. Int J Mol Sci. 2020;21(14).

40. (In preparation) Sophia K. Dunlap CAH, Mostafa Mohamed, Saeed Zakakhosravi, Taylor Schanel, Christopher D. Willey, Eun-Young Erin Ahn. Integrated transcriptomic and alternative splicing analyses reveal convergent biological programs in radiation-resistant glioblastoma.

41. Winer A, Adams S, Mignatti P. Matrix Metalloproteinase Inhibitors in Cancer Therapy: Turning Past Failures Into Future Successes. Mol Cancer Ther. 2018;17(6):1147–55.

42. Chandrashekar DS, Bashel B, Balasubramanya SAH, Creighton CJ, Ponce-Rodriguez I, Chakravarthi B, Varambally S. UALCAN: A Portal for Facilitating Tumor Subgroup Gene Expression and Survival Analyses. Neoplasia. 2017;19(8):649–58.

43. Chandrashekar DS, Karthikeyan SK, Korla PK, Patel H, Shovon AR, Athar M, Netto GJ, Qin ZS, Kumar S, Manne U, Creighton CJ, Varambally S. UALCAN: An update to the integrated cancer data analysis platform. Neoplasia. 2022;25:18–27.

